# STOCHASTIC MODELING OF HEMATOPOIETIC STEM CELL DYNAMICS

**DOI:** 10.1101/2025.01.27.635091

**Authors:** Carlos Alfaro-Quinde, Katerina E. Krstanovic, Paula A. Vásquez, Katie L. Kathrein

**Affiliations:** Department of Biological Sciences, University of South Carolina, Columbia, SC; Department of Biomedical Engineering, University of South Carolina, Columbia, SC; Department of Mathematics, University of South Carolina, Columbia, SC

## Abstract

The study of hematopoietic stem cell (HSCs) maintenance and differentiation to supply the hematopoietic system presents unique challenges, given the complex regulation of the process and the difficulty in observing cellular interactions in the stem cell niche. Quantitative methods and tools have emerged as valuable mechanisms to address this issue; however, the stochasticity of HSCs presents significant challenges for mathematical modeling, especially when bridging the gap between theoretical models and experimental validation. In this work, we have built a flexible and user-friendly stochastic dynamical and spatial model for long-term HSCs (LT-HSCs) and short-term HSCs (ST-HSCs) that captures experimentally observed cellular variability and heterogeneity. Our model implements the behavior of LT-HSCs and ST-HSCs and predicts their homeostatic dynamics. Furthermore, our model can be modified to explore various biological scenarios, such as stress-induced perturbations mediated by apoptosis, and successfully implement these conditions. Finally, the model incorporates spatial dynamics, simulating cell behavior in a 2D environment by combining Brownian motion with spatially graded parameters.

**\*Summary Statement:** This study addresses the challenge of characterizing hematopoietic stem cell (HSC) dynamics by developing a flexible, user-friendly stochastic spatial model of long-term and short-term HSCs. The model captures observed cellular variability and heterogeneity, predicts homeostatic dynamics, can be adapted to simulate stress-induced perturbations like apoptosis, and incorporates a spatial component to analyze HSC movement within a bone marrow niche.

## Introduction

Hematopoietic stem cells (HSCs) are highly dynamic yet rare cells residing within distinct hypoxic niches of the bone marrow microenvironment (Kiel & Morrison, 2006; Orkin & Zon, 2008). To preserve genomic integrity, the majority of HSCs remain in a quiescent state *in vivo*, limiting their exposure to cell-cycle-associated DNA damage (Pinho & Frenette, 2019; Pinho & Zhao, 2023; Seita & Weissman, 2010; Wilson et al., 2008). At the top of the hematopoietic hierarchy are long-term hematopoietic stem cells (LT-HSCs), which possess the unique ability to sustain long-term hematopoietic reconstitution when needed but also maintain the highest levels of quiescence as a protective mechanism. Upon activation, LT-HSCs differentiate into **s**hort-term hematopoietic stem cells (ST-HSCs), which exhibit reduced quiescence, yet are still capable of hematopoietic reconstitution. ST-HSCs subsequently give rise to the more heterogeneous pool of multipotent progenitor cells (MPPs), which contribute to the production of mature blood cells necessary for steady-state hematopoiesis and maintain very low levels of long-term reconstitution capabilities (Akashi et al., 2000; Benveniste et al., 2010; Kondo et al., 1997; Morrison et al., 1997; Yamamoto et al., 2013; L. Yang et al., 2005).

Regulation of HSCs is essential for maintaining hematopoietic homeostasis. HSCs have long been a central focus in stem cell biology due to their complex and dynamic processes, which are governed by both cell-intrinsic factors, such as epigenetic regulators and transcription factors, and cell-extrinsic signals from the bone marrow niche (Orkin & Zon, 2008; Pinho & Frenette, 2019; Wilson et al., 2008). These regulatory mechanisms ensure HSCs can generate all mature blood lineages as well as play a critical role in restoring the hematopoietic system following injury, including transplantation into marrow-ablated recipients and inflammatory conditions (Anderson et al., 2020; Baldridge et al., 2010; Jacobson et al., 1951; Porada et al., 2015; Thompson et al., 2024; Till & McCulloch, 1961; Walter et al., 2015; Wilson et al., 2008). Yet, the molecular mechanisms and underlying dynamics of HSC function remain incompletely understood (Paul et al., 2015). Emerging evidence indicates that HSCs make continuous contributions to hematopoiesis; however, the precise quantitative nature of these contributions remains under investigation. Experimental limitations, such as perturbations introduced during experimental data acquisition, often challenge the accurate assessment of HSC dynamics due to the disruption of the system.

Computational approaches such as mathematical modeling can represent critical components of biological systems without the need for experimental disruption (Bartocci & Lió, 2016; Fischer, 2008). By employing calculations and in-silico analyses, these models provide insights into how various factors influence system regulation, enabling predictions of complex biological behaviors. Such approaches not only guide future experimental research but also hold the potential to uncover novel therapeutic targets (Mackey & Maini, 2015).

Mathematical modeling has been instrumental in elucidating the contributions of HSCs to hematopoiesis. Early models, such as those proposed by Mackey,1978, and later refined by Dale & Mackey, 2015, have provided valuable insights into the dynamic transitions of HSCs between quiescence and proliferation. More recent models, such as the one by Kawahigashi et al., 2024, have examined age-related changes in the HSC pool, shedding light on the interplay among stem cell-stem cell (S-S) division, stem cell-progenitor cell (S-P) division, and progenitor cell-progenitor cell (P-P) division. Similarly, Bernitz et al., 2016, explored specific biological mechanisms of HSCs, proposing the existence of divisional history memory in HSCs. Mathematical models have also been utilized to investigate stress-induced HSC dynamics under both theoretical and experimental conditions. For instance, Dingli & Michor, 2006, and Parajdi et al., 2020, studied the behavior of leukemic HSCs within the hematopoietic system, while Bonnet et al., 2021, analyzed stress hematopoiesis in immature compartments, including LT-HSCs and ST-HSCs, over time in a murine model of phenylhydrazine (PHZ)-induced stress. These deterministic models provide valuable insights into the average HSC dynamics; however, they neglect the inherent stochasticity of HSC fates (Sussmann, 1978), a limitation our stochastic model directly addresses.

HSCs decision-making is inherently complex and not yet fully defined; however, the outcomes are well characterized: proliferation, differentiation, mobilization, or death. This complexity makes HSCs an ideal subject for investigation through stochastic modeling, which accounts for probabilistic events and individual cellular variability. Stochastic analyses have significantly enhanced our understanding of hematopoiesis. For instance, Xu et al., 2018, explored the hematopoietic system using stochastic simulations integrated within a graphical interface-mathematical modeling framework. Their work demonstrated the flexibility of stochastic simulations in testing hypotheses and exploring system dynamics. However, their compartmentalized design, which includes only two biological compartments—immature and mature cells—may oversimplify the system and overlook the distinct fates and dynamic behaviors of HSCs.

In addition to intrinsic factors, HSCs are closely regulated by their niche, which plays a pivotal role in their function, mobility, and overall maintenance. Recent mathematical models, such as the one proposed by Pedersen et al., 2023, have investigated the interactions between HSCs and their niche. Their model provides a valuable framework for analyzing how stem cell proliferation, differentiation, and attachment/detachment dynamics within the niche influence various clinical scenarios. While this model offers significant insights into the regulation of HSCs within the bone marrow microenvironment, it does not account for the stochastic movements and spatiotemporal dynamics of HSCs. Exploring these dynamics requires single-cell-level modeling that incorporates spatial data to describe HSC migration patterns. Such data enable the testing of hypotheses regarding the relative positions of different cell types, the mechanisms underlying HSC motility or immobility, and the factors driving these behaviors. Resolving HSC–niche dynamics comprehensively needs the integration of spatiotemporal data not only under homeostatic conditions but also under perturbations, both within individual niches and across the entire bone marrow environment.

To investigate the inherently variable dynamics of LT-HSCs and ST-HSCs, we developed a spatially explicit (2D) computational model. This model accounts for cellular heterogeneity and the probabilistic nature of HSC behavior. By fitting the model to two distinct *in vivo* experimental datasets describing the dynamics of phenotypically distinct LT-HSCs and ST-HSCs, we established a comprehensive theoretical configuration of homeostatic LT-HSCs and ST-HSCs in mice. The model accurately reproduces the observed experimental data and generates testable predictions. Moreover, we demonstrated the model’s flexibility by simulating apoptosis-induced stress scenarios in LT-HSCs and ST-HSCs. Finally, using the model’s spatial component, we explored the effects of varying gradients on quiescent and active LT-HSCs and ST-HSCs under homeostatic conditions, gaining insights into their spatial dynamics and localization.

## Results

We formulated a stochastic, agent-based model to simulate the behavior of long-term (LT-HSCs) and short-term (ST-HSCs) hematopoietic stem cells, incorporating key processes such as quiescence, apoptosis, distinct modes of cell division, and eventual inactivation after reaching a maximum number of divisions. Both LT-HSCs and ST-HSCs can enter a quiescent state with distinct probabilities. While quiescent, cells may either remain dormant or stochastically undergo apoptosis. Non-quiescent cells also face a probability of apoptosis. Surviving cells can divide via four distinct modes: symmetric self-renewal, asymmetric division, direct differentiation, or symmetric differentiation. The selection of each division mode is determined stochastically through a Markov process. This division process continues until LT-HSCs and ST-HSCs reach their respective maximum division limits, at which point they transition to an inactive state. Figure 1 provides a visual overview of the model; Supplementary Table 1 lists the parameters; and the Materials and Methods section provides further details, mathematical equations, and modeling assumptions.

**Figure 1.**
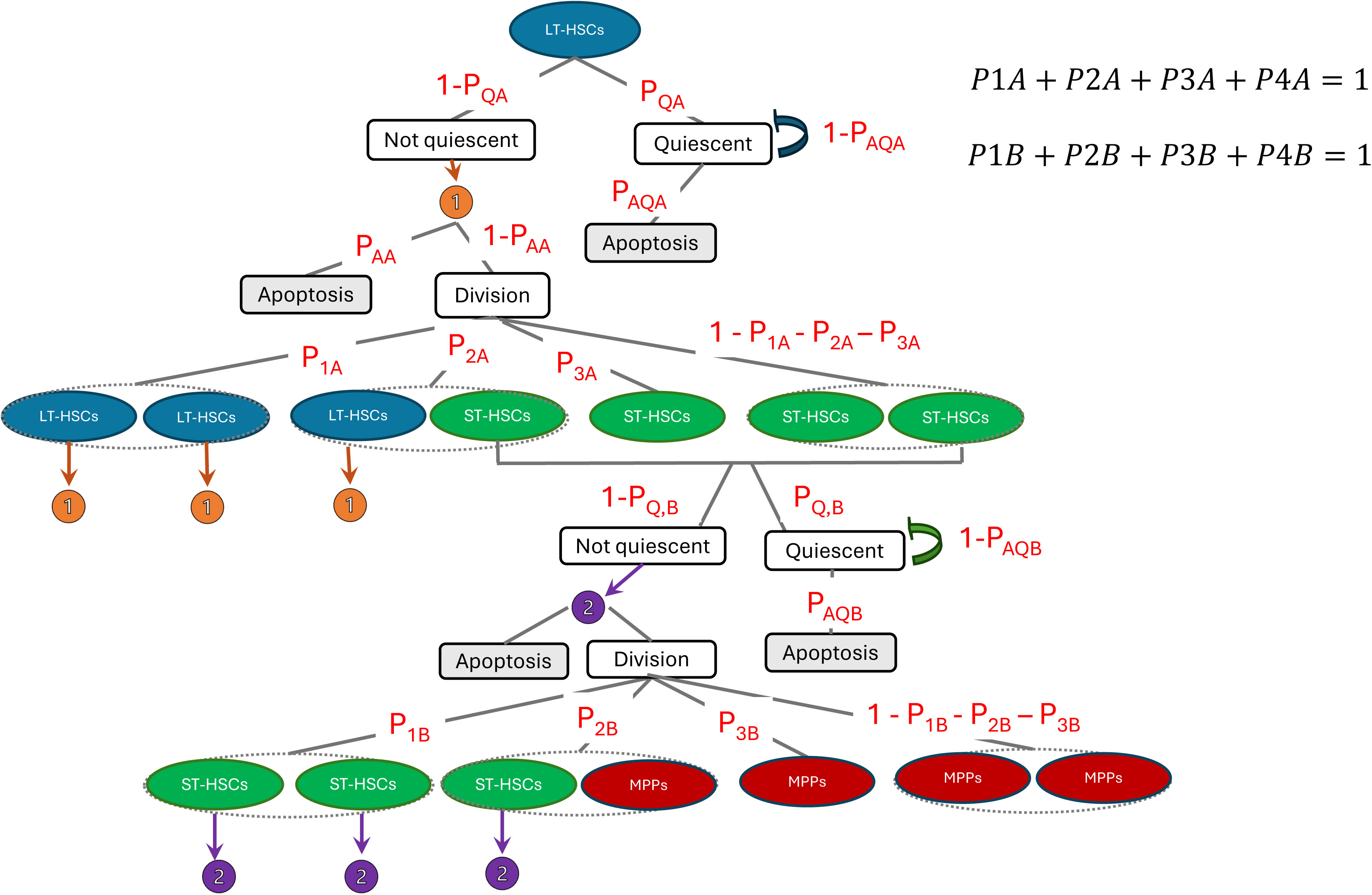
Schematic representation of the dynamical stochastic model of HSC dynamics. The model illustrates the dynamics of LT-HSCs, ST-HSCs, and MPPs. LT-HSC and ST-HSCs can be quiescent or divided based on PAA and PAB probabilities respectively. Both proliferating and dividing cells are susceptible to die based on PAA and PAB, and P*AQ*A and P*AQ*B respectively. Both LT-HSCs and ST-HSCs have 4 modes of division: symmetrical proliferation, asymmetrical proliferation, direct differentiation, and symmetrical differentiation. All four modes of division can contribute to the growth or decline of their respective population based on different probabilities. The arrows pointing down to numbers 1 and 2 indicate the direction of the model. LT-HSCs are depicted in green, ST-HSCs in blue, and MPPs are red.

### 1. Formulation of a homeostatic configuration of LT-HSCs and ST-HSCs

To obtain a model configuration that reflects homeostatic LT-HSCs and ST-HSCs, we fit our dynamical model to quantitative *in vivo* murine HSC data from two independent experimental studies: Busch et al., 2015, and Säwen et al., 2018. Both studies utilized distinct markers, Fgd5 and Tie2, to lineage trace the progression of phenotypically defined mice LT-HSC and ST-HSC populations over time. We assessed these two datasets independently to evaluate the efficacy and flexibility of our model and to account for the differences in the reported HSC populations. Busch et al. characterized LT-HSCs as a more quiescent and slowly differentiating population, whereas Säwen et al. described a more dynamic and actively dividing population that significantly contributes to downstream differentiation. By incorporating both datasets, we demonstrate the ability of our model to accommodate varying experimental observations and to capture the heterogeneity of LT-HSC and ST-HSC dynamics. By fitting the model to the experimental data, we estimated parameter values, which are presented for both case studies in Table 1. Figures 2A and 2B illustrate the homeostatic configurations predicted by the parameterized model over a 40-week period, with each figure corresponding to one of the two experimental references. Additionally, we extended our simulations to track the progression of each population (LT-HSCs and ST-HSCs) up to 80 weeks. To account for the inherent stochasticity of the system, 50 independent simulations were used in both cases, with error bars representing the variability across simulations. Our model reveals that the primary differences in LT-HSC behavior between the two datasets lie in their differentiation dynamics and ability to sustain the stem cell population over time. In the homeostatic configuration fit to Busch et al. data, LT-HSCs predominantly exhibit **s**ymmetrical self-renewal division (52%), with direct and symmetrical differentiation occurring at 30%. For ST-HSCs, the model shows a symmetrical division rate of 40%, and a direct and symmetrical differentiation rate of 45%, suggesting a more prominent differentiating ability role for this population (Table 1). The apoptosis rate was estimated at 1% for LT-HSCs and 2.5% for ST-HSCs per time step. Mean cell cycle durations were 5 ± 1.5 weeks for LT-HSCs and 3.5 ± 1.5 weeks for ST-HSCs, reflecting the inherent heterogeneity and stochasticity of division timing within each population. The maximum number of divisions was set at 2 for LT-HSCs and 3 for ST-HSCs before entering an inactive state. During the simulations, the proportion of actively participating LT-HSCs contributing to proliferation and differentiation was 36%, whereas 45% of ST-HSCs were actively engaged (Figure 2A).

**Figure 2.**
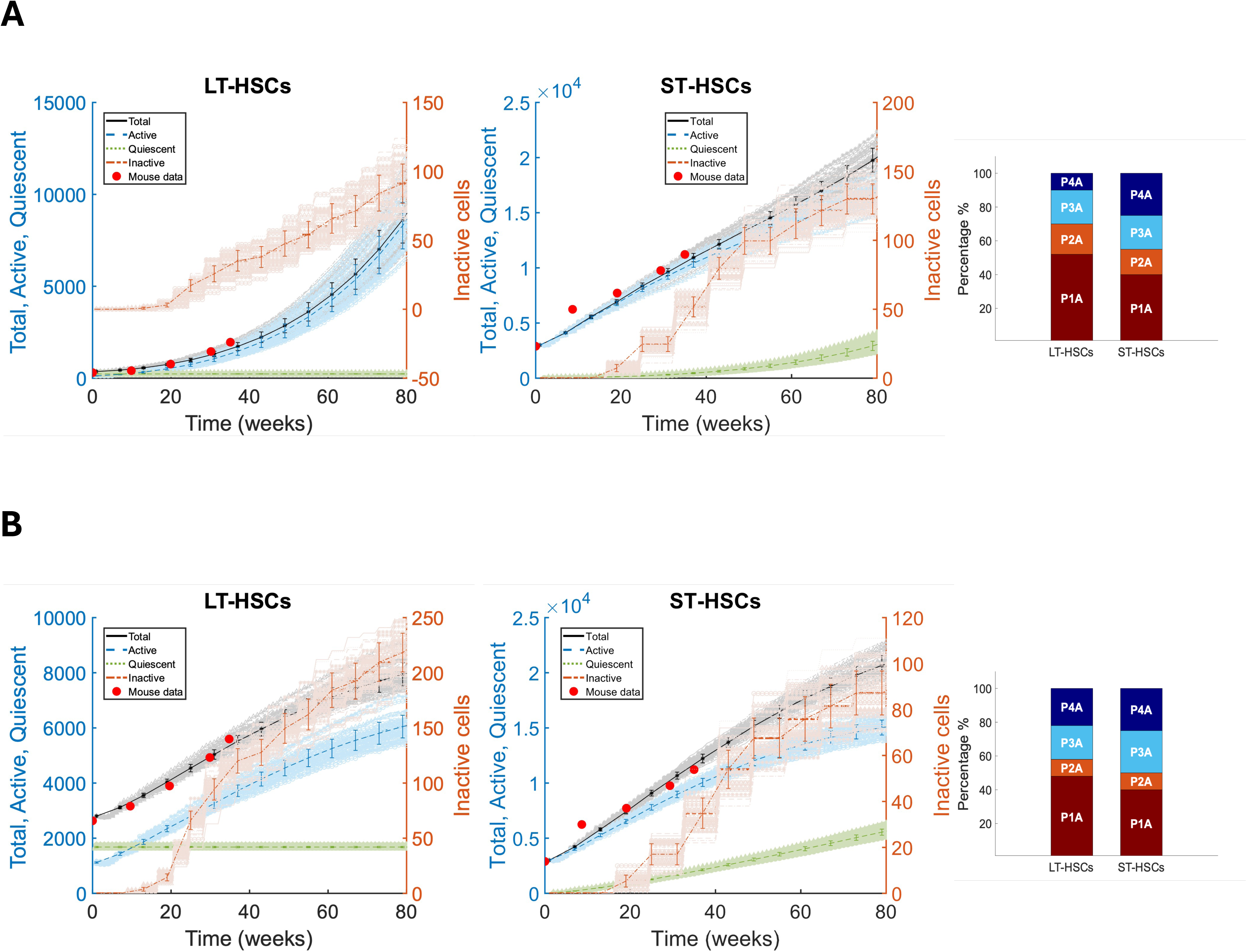
Homeostatic model simulations and predictions of LT-HSCs and ST-HSCs dynamics, based on Busch et and Säwen et al., up to 80 weeks. Figures A and B displayed the best fit of parameters for the dynamics of LT-HSCs and ST-HSCs, for a total of 80 weeks. The model is able to capture the dynamics proposed by **A)** Bush et al. and **B)** Säwen et al. and it is capable of predicting their future dynamics for another 40 weeks. Total cells (black), Active cells (blue), and Quiescent cells (green) are displayed on the right y-axis for both LT-HSCs and ST-HSCs, while the Inactive cells (orange) are displayed on the left y-axis. Experimental data from both independent datasets are shown as red dots. Solid lines represent the mean for each type of population, and the error bars are the SEM. seeds = 50. In addition, the division parameters used to fit the respective datasets are shown as stacked bar plots.

The homeostatic configuration of LT-HSCs and ST-HSCs fit to Säwen et al. data, also revealed a predominance of symmetrical self-renewal divisions in both populations, with rates of 48% for LT-HSCs and 40% for ST-HSCs. Additionally, the direct and symmetrical differentiation rates were higher compared to Busch et al., at 44% for LT-HSCs and 50% for ST-HSCs, suggesting that these populations are more actively engaged in differentiation during homeostasis. The proportion of actively participating LT-HSCs increased to 40% (compared to 36% in the Busch et al. configuration), while the percentage remained at 45% for ST-HSCs. This increased activation of LT-HSCs was accompanied by an elevated apoptosis rate of 2.5% for LT-HSCs, matching the rate observed for ST-HSCs. The mean cell cycle durations remained consistent with the Busch et al. configuration, at 5 weeks ± 1.5 weeks for LT-HSCs and 3.5 weeks ± 1.5 weeks for ST-HSCs (Table 1. Similarly, the maximum number of divisions before entering an inactive state remained unchanged, with LT-HSCs limited to 2 divisions and ST-HSCs to 3 divisions (Figure 2B).

Our model is able to simulate the homeostatic conditions with both experimental datasets, with Säwen et al showing the more accurate representation. This further highlights the importance of incorporating stochastic processes and individual cell fate variability in mathematical models to faithfully reflect the heterogeneity observed in LT-HSCs and ST-HSCs populations.

### 2. Modeling mice LT-HSCs and ST-HSCs stress response

To investigate the impact of perturbations within our model system, we examined how elevated inflammation— and the associated apoptosis induced by excessive pro-inflammatory cytokines during CAR-T cell therapy— affects the dynamics of LT-HSCs and ST-HSCs. This analysis was guided by the findings of Read et al, 2023

CAR-T cell therapy has emerged as a groundbreaking treatment for relapsed hematological cancers. However, the clinical application can result in significant adverse effects, most notably cytokine release syndrome (CRS), which arises from a surge of pro-inflammatory cytokines following T-cell activation. Therefore, patients often experience complications such as cytopenia and delayed bone marrow recovery (Fried et al., 2019; Schuster et al., 2019; Sermer & Brentjens, 2019). Read et al., 2023 investigated the effects of CAR-T cell therapy in a murine tumor model treated with human CAR-T cells. Their findings revealed that the mice developed CRS, characterized by elevated expression of pro-inflammatory cytokines. Notably, they observed that hematopoietic stem and progenitor cells (HSPCs) exhibited **i**ncreased apoptosis following CAR-T cell therapy, leading to a reduction in actively dividing LT-HSCs, while the quiescent LT-HSCs and ST-HSC populations remained unaffected. Furthermore, they demonstrated that the injection of pro-inflammatory cytokines into mice resulted in an increased Ki67% in ST-HSCs, indicating that a greater proportion of these cells exited quiescence during apoptosis-inducing inflammation.

To address the impact of apoptosis-related perturbations within our model system, we incorporated the following assumptions: i) Homeostatic conditions are based on the LT-HSC and ST-HSC configuration described by Säwen et al., 2018 ii) Quiescent LT-HSCs and ST-HSCs remain unaffected, consistent with their experimental findings. iii) Apoptosis rates for both LT-HSCs and ST-HSCs remain constant throughout the simulation.

Then, to study the effects of apoptosis-related changes over time, we considered three distinct scenarios based on the experimental data reported by Read et al., 2023 a) Increased apoptotic levels of LT-HSCs. b) Increased apoptotic levels of both LT-HSCs and ST-HSCs, accompanied by an increase in actively dividing ST-HSCs. c) Modification of the ST-HSC configuration to restore homeostasis under inflammatory conditions.

We mimicked these scenarios by increasing apoptosis rates for LT-HSCs and ST-HSCs and adjusting the homeostatic configuration of ST-HSCs to align with the Säwen et al., 2018 homeostatic values. The list of parameters for each scenario is provided in Table 2. To account for stochasticity, we ran 50 independent simulations for each scenario, with each modeled up to 100 weeks.

a. For the first scenario, we increased the apoptotic rate of LT-HSCs to 5% and 10% per time step, as experimentally reported by Read et al., while keeping the ST-HSC homeostatic configuration unchanged. Our model demonstrates the detrimental effects of elevated apoptosis on the LT-HSC population. Additionally, the simulation reveals a corresponding decrease in the ST-HSC population, likely due to the direct feedback dependency of ST-HSCs on LT-HSCs. However, when the LT-HSC apoptosis rate was increased to 5% (indicated in red), the ST-HSCs were able to partially recapitulate the homeostatic configuration proposed by Säwen et al., suggesting a compensatory role of ST-HSCs in the presence of non-functional LT-HSCs (Figure 3A).
b. For the second scenario, we increased the apoptosis rate of ST-HSCs to 10%, while maintaining the LT-HSC apoptosis rate at 5%. No changes were made to the homeostatic configuration of the LT-HSCs. To reflect the experimentally observed increased Ki67% reported for ST-HSCs by Read et al., we elevated the percentage of actively participating ST-HSCs to 70%. Our model demonstrates the detrimental impact of elevated apoptosis on LT-HSCs, leading to a reduction in the number of active LT-HSCs over time without completely depleting the population within the 100-week simulation timeframe (Figure 3B) Furthermore, the results indicate that increasing the proportion of actively participating ST-HSCs to 70% is insufficient to recapitulate the homeostatic dynamics reported by Säwen et al. (Figure 3B).
c. For the third scenario, we modeled the theoretical biological requirements for ST-HSCs to achieve homeostasis under conditions of apoptosis-related inflammation affecting both LT-HSCs and ST-HSCs. The apoptotic rate was set to 5% for LT-HSCs and 10% for ST-HSCs, while the percentage of actively participating ST-HSCs was increased to 70%, as in the second scenario. Our model demonstrates that for ST-HSCs to **r**ecapitulate homeostasis proposed by Säwen et al., the rate of symmetrical self-renewal division must increase to 70%, while direct and symmetrical differentiation is maintained at 20%. Under these conditions, the ST-HSCs successfully restored homeostatic dynamics for up to 40 weeks. However, beyond this time point, the ST-HSC population exhibited a linear decline, ultimately failing to sustain homeostasis for the remainder of the simulation (Figure 3C).

**Figure 3.**
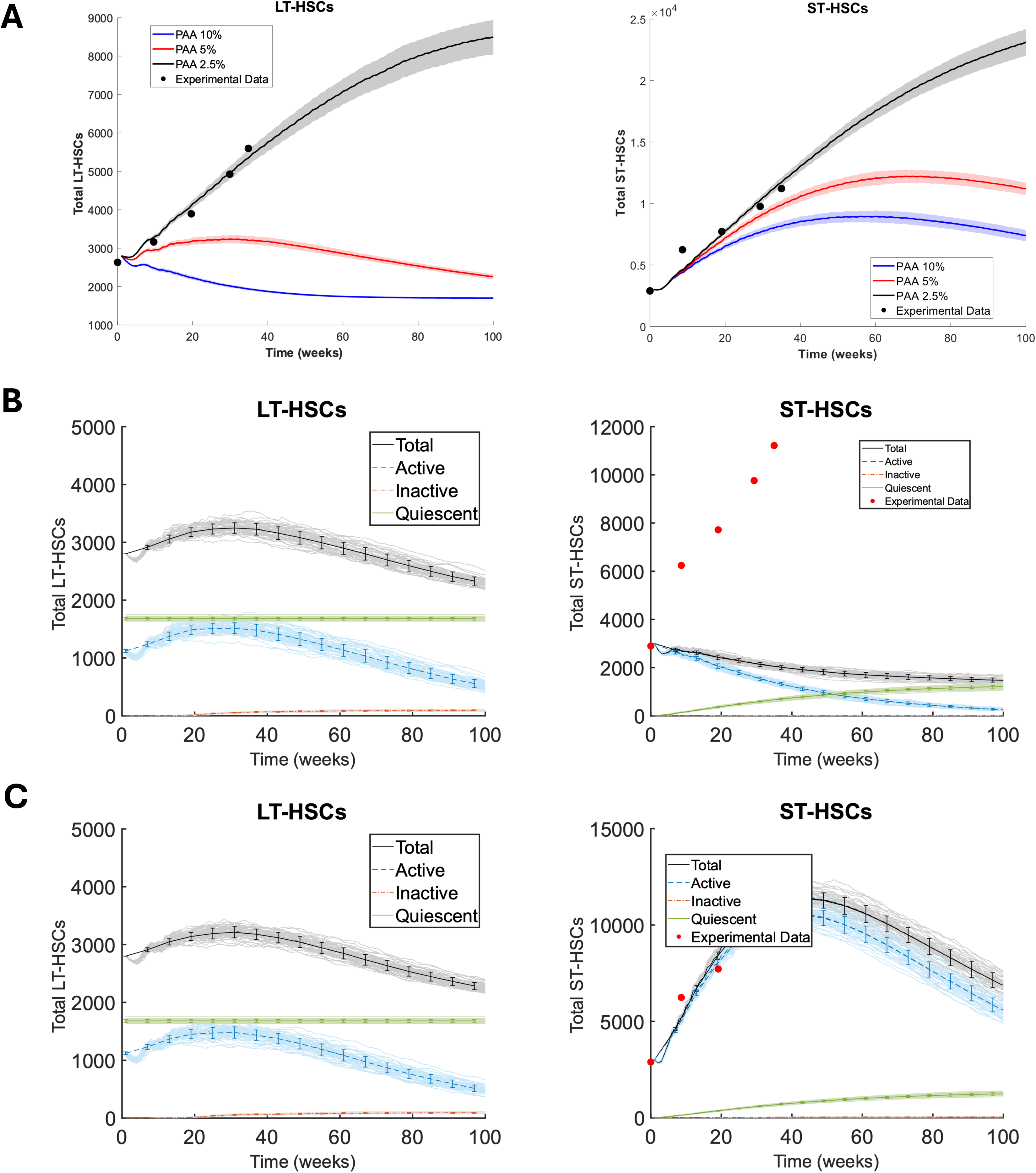
Theoretical apoptotic-inflammatory stress scenarios of LT-HSCs and ST-HSCs dynamics. **Figure A** displays the behavior of LT-HSCs under apoptotic stress conditions suggested by Read et al., after exposition to pro-inflammatory cytokine, up to 100 weeks. Three apoptotic rates are displayed: *PAA*=2.5% (black), *PAA*= 5% (red), and *PAA*=10% (blue). Then the effect of the three different LT-HSCs apoptosis rates on ST-HSCs dynamics is displayed while the homeostatic configuration of ST-HSCs was not modified. The experimental data from Säwen et al., is shown as black dots up to 40 weeks. Solid lines represent the mean for each type of population, and the shaded area is SEM. seeds = 50. **Figure B** shows LT-HSCs dynamics when *PAA* is modified to 5%. Behavior and ST-HSCs dynamics when *PAB* is modified to 10% and *PQB* is 0.3. The x-axis displays the time frame of the simulation, and the y-axis represents the population: Total cells (black), Active cells (blue), Quiescent cells (green), and Inactive cells (orange). The experimental data from Säwén et al., is shown as black dots up to 40 weeks. Solid lines represent the mean for each type of population, and the shaded area is SEM. seeds = 50. **Figure C** displays apoptotic-stress LT-HSCs dynamics when *PAA* is modified to 5%. In addition, ST-HSCs dynamic is displayed when *PAB* is modified to 10%, PQB=0.3, *P1B=0.7, P2B=0, P3B=0.1, P4B=0.2*. The x-axis displays the time frame of the simulation, and the y-axis represents the population: Total cells (black), Active cells (blue), Quiescent cells (green), and Inactive cells (orange). The experimental data from Säwén et al., is shown as black dots up to 40 weeks. Solid lines represent the mean for each type of population, and the shaded area is SEM. seeds = 50

### 3. Parameter Sensitivity analysis

We conducted a Latin Hypercube Sampling with Partial Rank Coefficient Analysis (LHS/PRCC) to evaluate the impact of various model parameters on the total number of active, quiescent, and inactive LT-HSCs and ST-HSCs. This analysis is essential for identifying the parameters that exert the strongest influence on the model outputs, thereby assessing the robustness of the dynamical model predictions.

For the active LT-HSCs, PRCC analysis shows *P1A, P2A,* and *divA* having a significant positive correlation with the output. On the other hand, *P3A, meanCCA, PQA,* and *PAA* exhibited negative correlations, with *PAA* displaying the highest significant negative relationship (Figure 4A). For the active ST-HSCs, PRCC identified positive correlations for *P2A, P1B, P2B*, and *divB*. Conversely, *P3A, P3B, PQA, PQB, PAA, PAB, meanCCA*, and *meanCCB* were negatively correlated, with *PAA* standing out as the most negatively statistically significant parameter, followed by *PAB* (Figure 5A).

**Figure 4.**
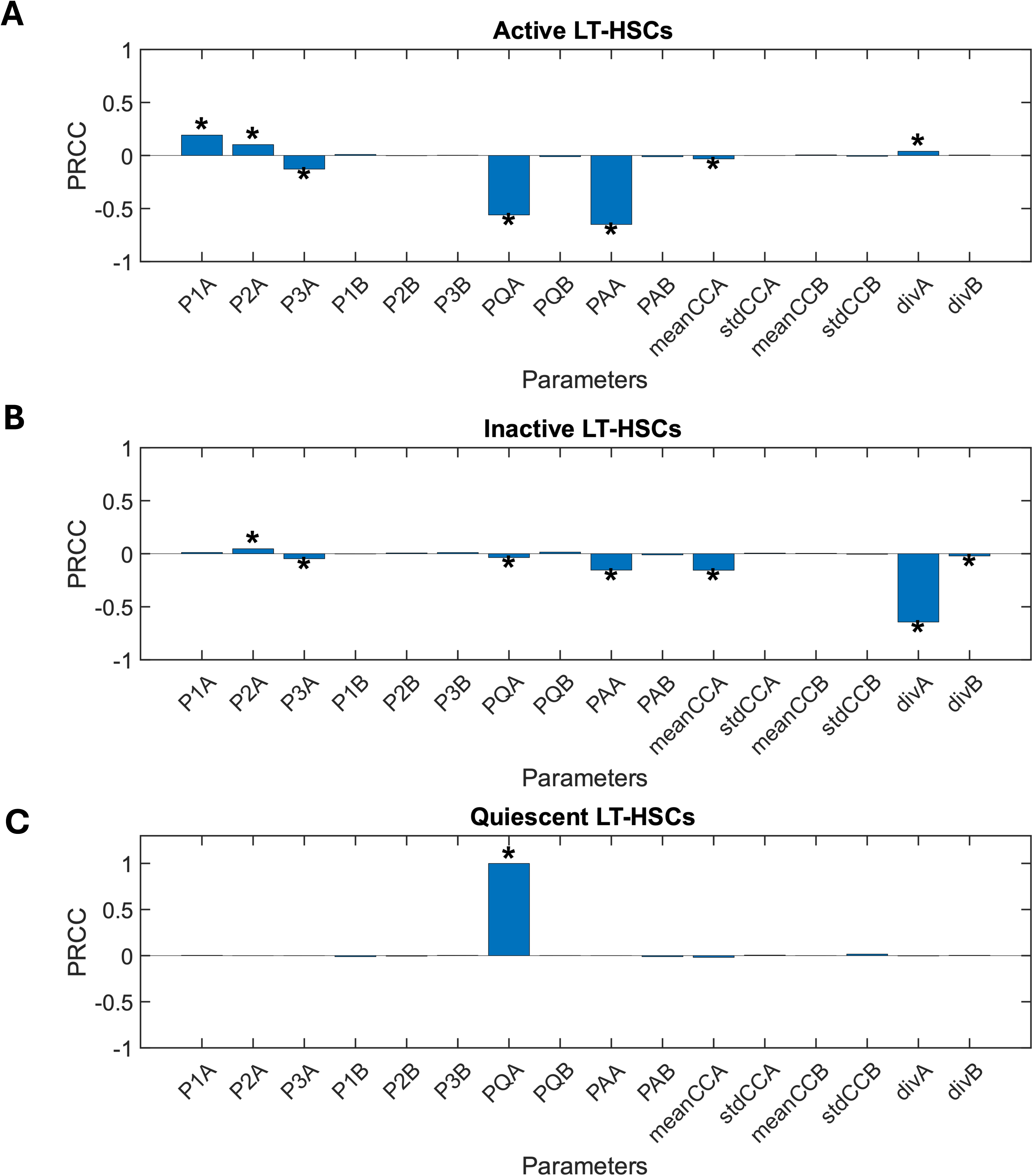
Partial Rank Coefficients (PRCC) of the total number of active, inactive, and quiescent LT-HSCs. For each parameter, the absolute value of its PRCC represents the sensitivity of the parameter-the larger the value is, the more sensitive the total number of **A)** active LT-HSCs, **B)** inactive LT-HSCs and **C)** Quiescent LT-HSCs, are to the corresponding parameter. The number of combinations = 10337. *Represents the value of PRCC, with a significance value <0.05.

**Figure 5.**
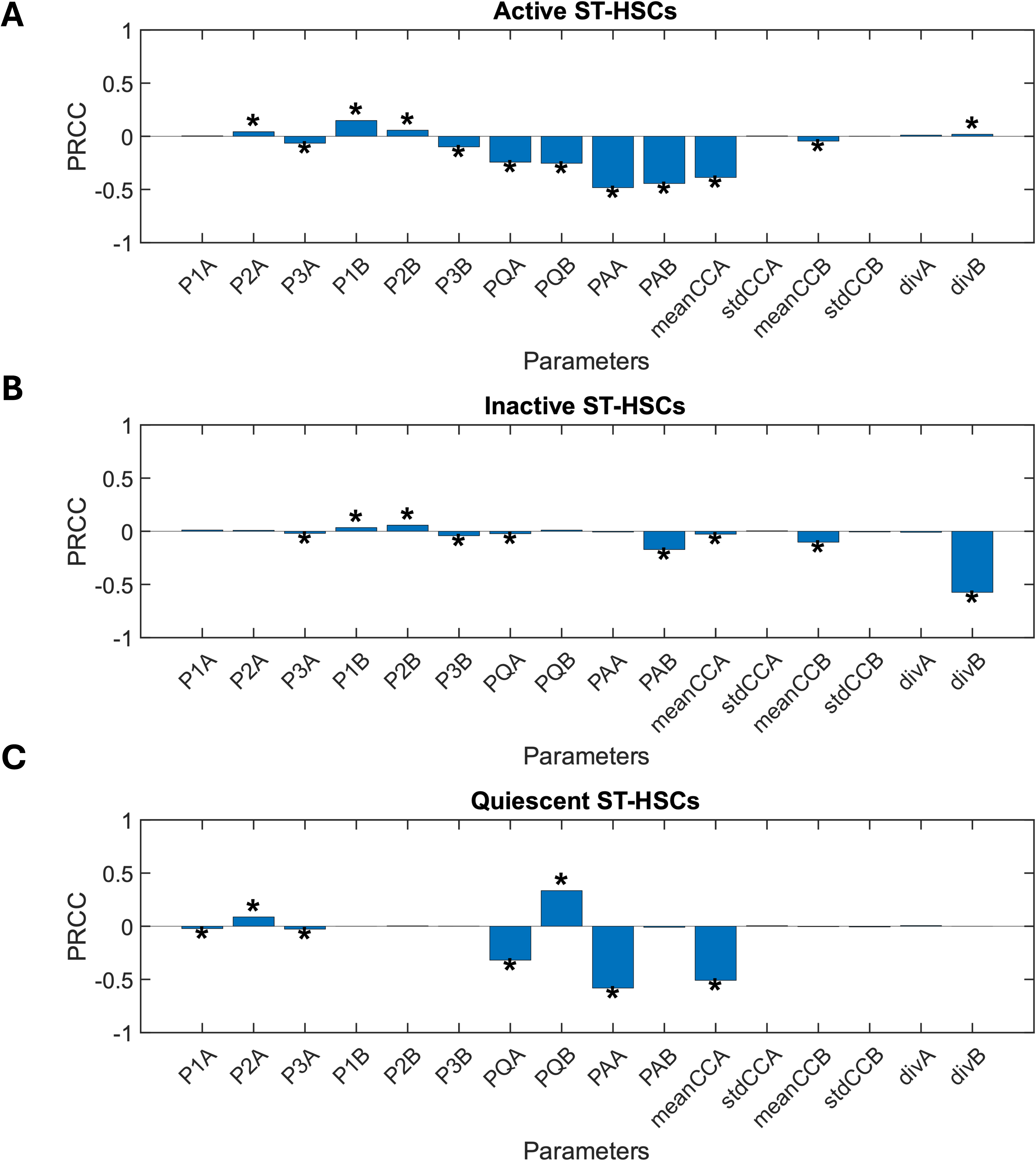
Partial Rank Coefficient (PRCC) of the total number of active, inactive and quiescent ST-HSCs with respect to the dynamical parameters. For each parameter, the absolute value of its PRCC represents the sensitivity of the parameter-the larger the value is, the more sensitive the total number of **A)** active ST-HSCs, **B)** inactive ST-HSCs and **C)** Quiescent ST-HSCs, are to the corresponding parameter. The number of combinations = 10337. * Represents the value of PRCC, which is not zero significantly, with a significance value <0.05.

For inactive LT-HSCs, negative correlations were found for *P3A, PQA, PAA*, *meanCCA, divB*, and *divA*, with the former one emerging as the parameter with the most significant negative influence on the inactive LT-HSCs (Figure 4B). Among inactive ST-HSCs, *P1B* and *P2B* demonstrated significantly positive relationships with the population size. In contrast, parameters such as *P3A, P3B, PQA, PAB, meanCCA, meanCCB,* and *divB* had significant negative correlations, with *divB* showing the strongest negative influence (Figure 5B).

For quiescent LT-HSCs, PRCC analysis revealed that *PQA* was the only significant parameter displaying a positive correlation, with a PRCC value of 1 (Figure 4C) Meanwhile, for quiescent ST-HSCs, positive correlations were observed for *P2A* and *PQB*. However, parameters like *P1A, P3A, PQA, PAA*, and *meanCCA* were negatively correlated with the output (Figure 5C).

### 4. Exploring the Influence of Spatial Gradients on LT-HSC and ST-HSC Dynamics

Using the Euler-Maruyama method for stochastic integration, our model simulates the Brownian motion of active and quiescent LT-HSCs and ST-HSCs within a two-dimensional space representing a single bone marrow niche. The spatial model incorporates two sets of parameters: (1) spatial parameters, including niche size (x and y dimensions), diffusion coefficient, and spatial gradients for the probabilities of quiescence (*pAQ* and *pBQ*) and mean cell cycle duration (*meanCCA* and *meanCCB*) (Supplementary Table 1); and (2) parameters defining the homeostatic configuration, fitted to experimental data published by Säwen et al., 2018. All model parameters are summarized in Table 3.

To explore the spatial dynamics of LT-HSCs and ST-HSCs, we systematically varied *aQ* (representing the gradient in the probability of a cell becoming quiescent, *pAQ* or *pBQ*) and *aD* (representing the gradient in the mean proliferation rate, *meanCCA* or *meanCCB*) to create three numerical combinations representing high, medium, and low spatial gradients (see Materials and Methods). Our numerical analysis demonstrates the formation of significant spatial gradients for both active and quiescent LT-HSCs and ST-HSCs as the parameters *aD* and *aQ* are varied. These gradients highlight the sensitivity of cell distribution and heterogeneity to spatial cues, providing insights into the dynamic behavior of stem cells within the niche.

Figure 6A illustrates the spatial distribution of active LT-HSCs under different gradient conditions. Under high gradient conditions (*aQ* = 0.25, *aD* = 0.25), the initial uniform distribution of cells at time step 20 gives way to a gradual accumulation of cells in the top-left corner of the domain by time step 80. This clustering pattern is likely driven by the combined effects of the increasing probability of quiescence from left to right and the decreasing mean division time from bottom to top. With a medium gradient (*aQ* = 0.5, *aD* = 0.5), the clustering is less pronounced, resulting in a broader distribution of cells across the domain. Under low gradient conditions (*aQ* = 0.75, *aD* = 0.75), where the parameters vary minimally across the domain, cells remain relatively uniformly distributed throughout the simulation.

**Figure 6.**
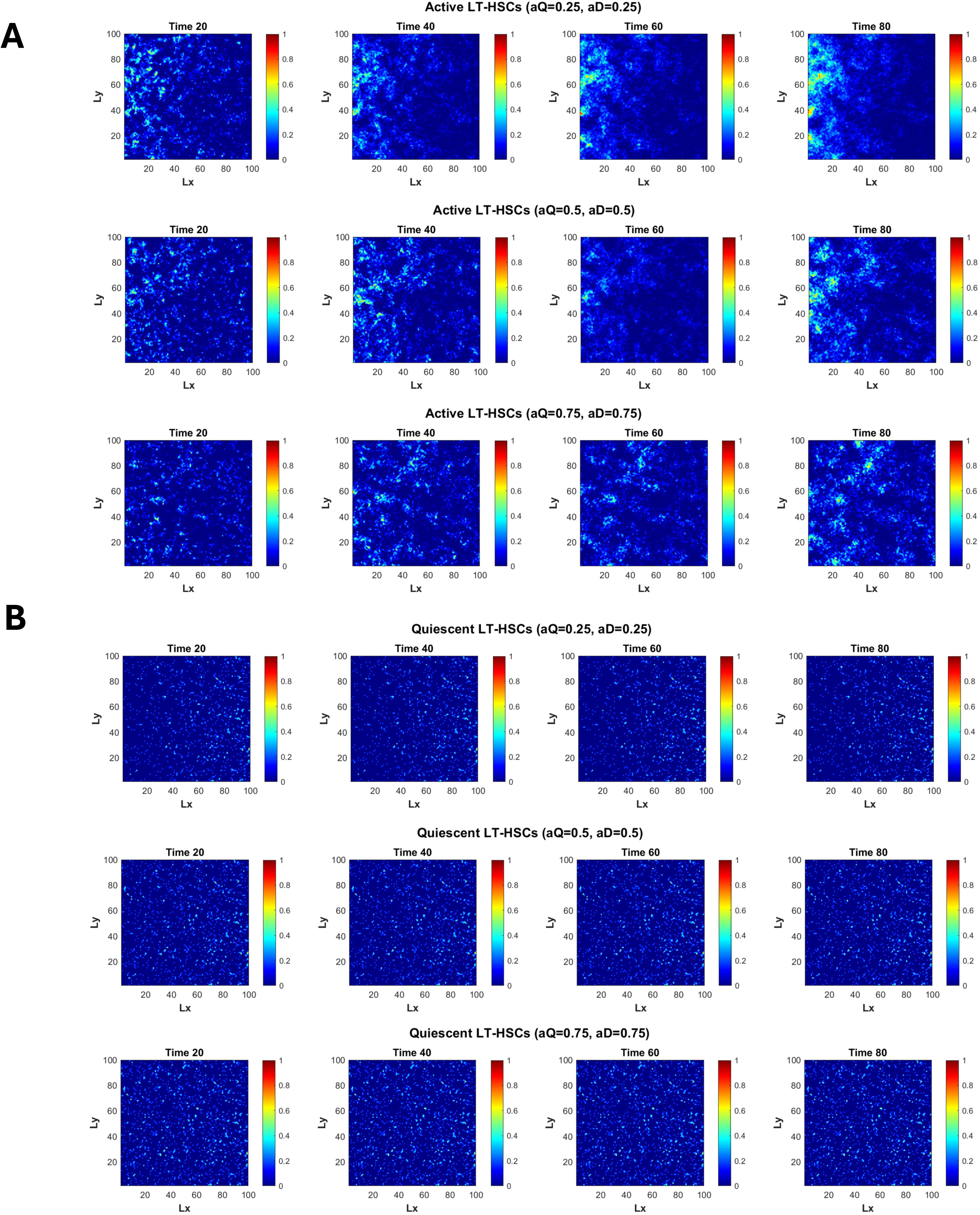
Theoretical numerical simulations of active and quiescent LT-HSCs under different *aQ* and aD gradient parameters. The figure displays the spatial distribution of active and quiescent LT-HSCs within the 2-D dimensional space, mimicking the bone marrow niche, under a range of uniform *aQ* and aD combinations = 0.25, 0.5 and 0.75 at time steps: 20, 40, 60 and 80. We used the homeostatic configuration used to fit Säwén et al., dataset. Spatial distribution is shown as a 2-D color histogram where 1 means the highest density and 0 is the lowest density. The x-axis corresponds to the L_x_ and the y-axis is the L_y_ spatial boundaries. **A)** Active LT-HSCs when *aQ* and *aD* = 0.25, *aQ* and *aD* = 0.5, and *aQ* and *aD* = 0.75, and **B)** Quiescent LT-HSCs when *aQ* and *aD* = 0.25, *aQ* and *aD* = 0.5, and *aQ* and *aD* = 0.75.

Similar spatial distribution patterns are observed for active ST-HSCs, as shown in Figure 7A. Under high gradient conditions, ST-HSCs also accumulate in the top-left corner by time step 80. With medium gradients, the clustering is less pronounced, and under low gradients, the distribution remains relatively uniform.

**Figure 7.**
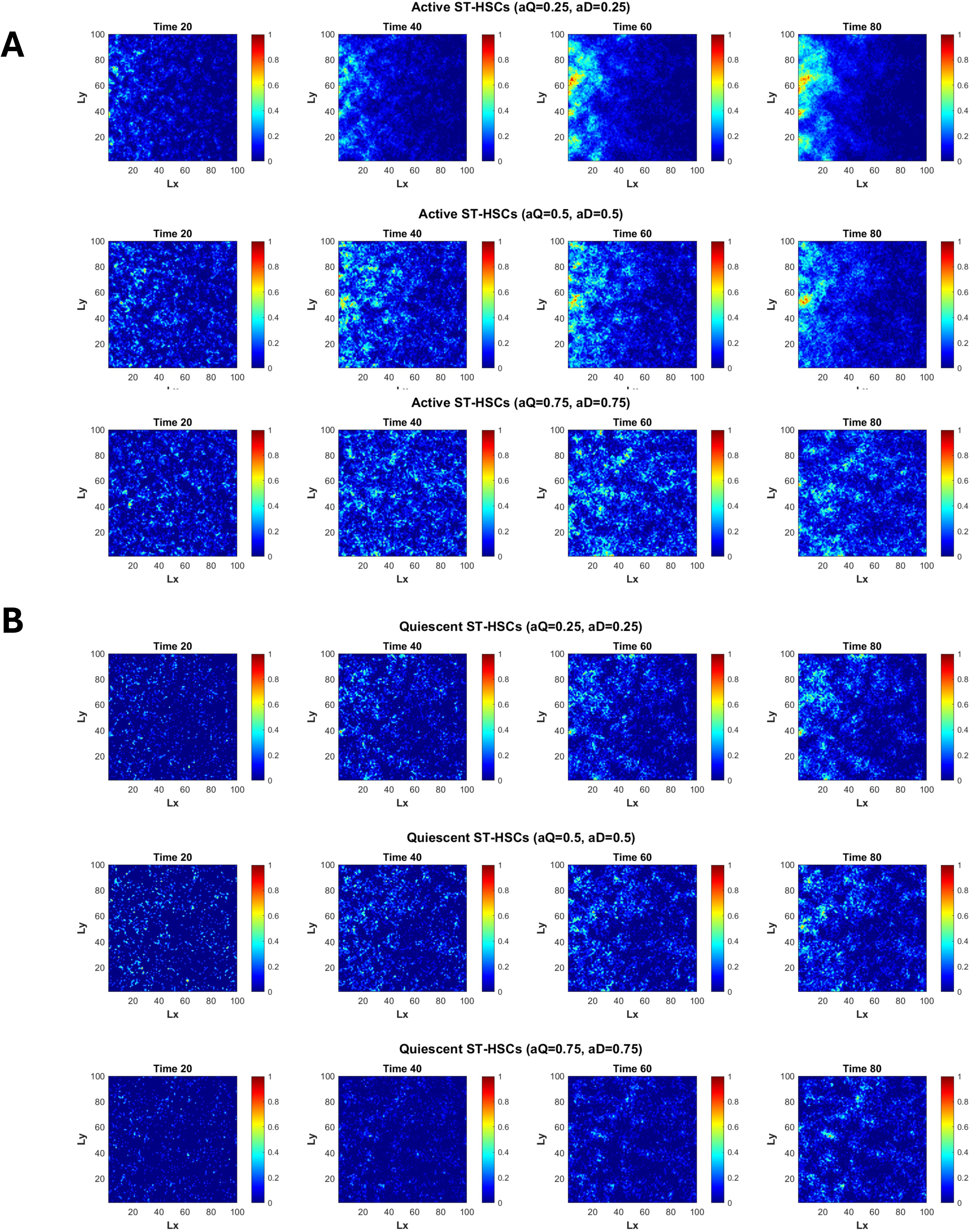
Theoretical numerical simulations of active and quiescent ST-HSCs under different *aQ* and aD gradient parameters. The figure displays the spatial distribution of active and quiescent ST-HSCs within the 2-D dimensional space, mimicking the bone marrow niche, under a range of uniform *aQ* and *aD* combinations = 0.25, 0.5 and 0.75 at time steps: 20, 40, 60 and 80. We used the homeostatic configuration used to fit Säwén et al., dataset. Spatial distribution is shown as a 2-D color histogram where 1 means the highest density and 0 is the lowest density. The x-axis corresponds to the L_x_ and the y-axis is the L_y_ spatial boundaries. **A)** Active ST-HSCs when *aQ* and *aD* = 0.25, *aQ* and *aD* = 0.5, and *aQ* and *aD* = 0.75, and **B)** Quiescent ST-HSCs when *aQ* and *aD* = 0.25, *aQ* and *aD* = 0.5, and *aQ* and *aD* = 0.75.

For quiescent LT-HSCs under high gradient change (*aQ*=0.25, *aD*=0.25), the cells were initially uniformly distributed across the domain at time step 20, with a slight concentration observed on the right side. As time progressed (time steps 40, 60, and 80), the cells increasingly clustered on the right side, with some spreading towards the left side, where most of the active LT-HSCs are located. Under medium gradient (*aQ*=0.5, *aD*=0.5), the quiescent LT-HSCs displayed a random distribution across the entire spatial domain. This pattern remained largely unchanged over time, with only minor gradient adjustments observed by time step 80. In the low-gradient scenario (*aQ*=0.75, *aD*=0.75), the quiescent LT-HSCs were even more distributed throughout the spatial domain. No clear clustering patterns emerged during any of the time steps (20, 40, 60, or 80), reflecting the limited or low variability of the parameters across the spatial domain (Figure 6B).

Finally, for quiescent ST-HSCs, the behavior in terms of gradient and cell location was similar across all levels (high, medium, and low), with the primary distinction being the increased number of cells throughout the simulation. Across all time steps and gradient change levels, no clear clustering patterns were observed, as the cells appeared to remain uniformly distributed across the spatial domain (Figure 7B).

## Discussion

The intricate dynamics between LT-HSCs and ST-HSCs, particularly their self-renewal and differentiation rates, play a critical role in maintaining hematopoietic homeostasis (Mann et al., 2022). Disruptions in this balance can lead to stress hematopoiesis. While previous studies have employed ODEs to quantitatively model HSC behavior, such approaches often pose interpretability challenges and fail to account for the **s**tochasticity and intrinsic heterogeneity inherent in HSC populations.

In this work, we present a flexible stochastic model of LT-HSC and ST-HSC behavior. The model incorporates a two-dimensional spatial motion component to simulate the dynamics of quiescent and active LT-HSCs over time, providing a robust framework for studying HSC behavior and interactions in both normal and perturbed conditions.

### Homeostatic Configuration of LT-HSCs and ST-HSCs

To validate our model, we fit *in vivo* label-tracing data from two independent homeostatic studies, each describing distinct phenotypic behaviors of LT-HSCs and ST-HSCs. Our model successfully captured the stochastic dynamics of LT-HSCs and ST-HSCs under both scenarios and revealed key differences in LT-HSC and ST-HSC dynamics between the two datasets, particularly in differentiation and population maintenance. In the homeostatic configuration fitted to the Busch et al. data (Figure 2A), the proportions of actively participating LT-HSCs and ST-HSCs were 36% and 45%, respectively, aligning with their experimental observations that at least 30% of HSCs contribute to hematopoiesis during homeostasis, including symmetrical divisions and differentiation (Busch et al., 2015). In addition, we showed that symmetrical division rates are predominant over the simulation, with 52% for LT-HSCs and 48% for ST-HSCs, consistent with the findings by Kawahigashi et al., 2024, and Loeffler et al., 2019. Moreover, the combined differentiation rates were 30% for LT-HSCs and 44% for ST-HSCs, while LT-HSC apoptosis was 1% per time step, as previously reported by Foudi et al., 2009. Our simulation ultimately led to an exponential increase in LT-HSCs and a linear increase in ST-HSCs over 80 weeks. While the simulated exponential growth of LT-HSCs is not biologically relevant, during homeostatic conditions, it likely reflects the low apoptosis rate estimated from the data, consistent with the expected quiescence of LT-HSCs. Our best model fit using the Säwen et al. dataset (Figure 2B), shows LT-HSC activity increased to 40%, with ST-HSC participation remaining at 45%. Symmetrical division rates were 48% for LT-HSCs and 40% for ST-HSCs, while the total differentiation rates were 44% for LT-HSCs and 50% for ST-HSCs. Apoptosis rates for both cell types increased to 2.5%, as reported by Barile et al., 2020, resulting in linear growth of LT-HSC and ST-HSC populations, consistent with Kawahigashi et al., 2024 findings. Our simulations suggest a more dynamic HSC phenotypic profile with increased differentiation and activation compared to the Busch et al. dataset.

A central assumption of our model is the division potential of HSCs. Based on Bernitz et al., 2016, we proposed that LT-HSCs and ST-HSCs enter an inactive state after 2 and 3 divisions, respectively. Our simulations showed that inactive cell numbers remained small compared to active cells, emphasizing that homeostasis is driven by the percentage of active cells rather than by the ability of individual HSCs to divide multiple times. These findings demonstrate the utility of our model in uncovering heterogeneity in HSC behavior and providing insights into their dynamics under homeostatic conditions.

### Modeling apoptotic-stress conditions on LT-HSCs and ST-HSCs

To further validate our model, we examined the effects of apoptosis-related stress on the homeostatic configuration of LT-HSCs and ST-HSCs under three distinct scenarios, using experimental data from Read et al., 2023 to mimic apoptotic stress in our model. In the first scenario, we found that increased LT-HSC apoptosis rates of 5% and 10% (compared to the homeostatic rate of 2.5%) showed significant depletion of the active LT-HSC pool (Figure 3A), with mild and severe depletion respectively, aligning with findings by (Read et al., 2023; Reagan & Rosen, 2015). Quiescent LT-HSCs and ST-HSCs remained unaffected due to unchanged quiescent apoptosis parameters (*PAQA, PAQB*), consistent with Read et al., 2023. Therefore, no complete depletion of the total number of LT-HSCs was observed during the simulations. 5% LT-HSC apoptosis reduced ST-HSC numbers but allowed partial compensation, maintaining homeostasis for up to 30 weeks as ST-HSCs increased their contribution to differentiation, aligning with findings from Säwen et al., 2018 and other studies on ST-HSC adaptability under stress (Bhattacharya et al., 2006; Yamamoto et al., 2013).

In the second scenario, fixing apoptotic rates at 5% for LT-HSCs and 10% for ST-HSCs showed declines in both active LT-HSCs and ST-HSCs. The active LT-HSCs were almost depleted towards the end of the simulation (Figure 3B), while ST-HSC depletion also showed the direct elevated apoptosis and feedback from LT-HSCs. Increasing ST-HSC symmetrical proliferation (*P1B*) to 70% allowed a compensatory increase in proliferative ST-HSCs, consistent with Read et al., 2023 and Singh et al., 2020, but failed to fully recapitulate homeostasis, suggesting that changes in self-renewal and differentiation capacities are required for ST-HSCs to sustain homeostasis under stress conditions.

In the third scenario, we mimicked experimental homeostasis data under inflammatory conditions (Säwen et al., 2018) for ST-HSCs. Symmetrical self-renewal was increased to 70%, with symmetric differentiation at 20%, enabling ST-HSCs to maintain downstream compartments despite apoptosis-induced inflammation (Barile et al., 2020). While this adjustment maintained homeostasis for up to 40 weeks, ST-HSC numbers declined linearly beyond this period, suggesting that quiescent cells failing to exit G0 under stress may lead to delayed recovery and eventual depletion of these compartments.

Our findings emphasize the intricate interplay between apoptotic stress and HSC dynamics, shedding light on the processes governing the balance between apoptosis, self-renewal, and differentiation required to sustain hematopoietic homeostasis under stress. By aligning with experimental observations, we demonstrate that our model offers a robust framework for exploring the stochastic dynamics of LT-HSCs and ST-HSCs. Furthermore, modifying the model parameters enables the simulation and investigation of diverse stress-induced hematopoietic dysfunction scenarios over time, providing valuable insights into the underlying mechanisms driving these processes.

### Parameter Sensitivity analysis

PRCC analysis showed the sensitivity of LT-HSC and ST-HSC populations to key parameters (Figs. 4A–5C). For active LT-HSCs, *PQA* and *PAA* were the most significant parameters. Increasing *PQA* would lead to increased quiescent cells, decreasing the number of active LT-HSCs, while increasing *PAA* would reduce the production of active LT-HSCs due to elevated apoptosis. Similarly, for the active ST-HSCs, *PQA* and *PAA* negatively impacted the population, due to the direct feedback LT-HSCs have on ST-HSCs. Among all the significant parameters (Figure 5A), *PAA* with *PAB* had the greatest PRCC values impacting negatively the growth of active ST-HSCs, because increasing these parameters would lead to increased apoptosis.

For inactive LT-HSCs, the most influential and significant parameter was the *divA* (Figure 4B). Increasing *divA* would lead to a higher threshold the active LT-HSCs need to reach in order to enter into an inactive state. For the inactive ST-HSCs, various parameters were significant (Figure 5B), being the result of the direct feedback from the LT-HSCs. In addition, the most influential parameter was *divB* (PRCC ≈ -0.6), which controls the number of divisions ST-HSCs need to perform to enter into an inactive state.

For quiescent LT-HSCs, *PQA* was the sole positively influential and correlated parameter (PRCC = 1), aligning with model assumptions and behavior (Figure 5C). For quiescent ST-HSCs, *P2A* and *PQB* had positive effects, while *PQA*, *PAA*, and *meanCCA* negatively impacted their population. *PAA*, (PRCC ≈ -0.6), was the most significant, highlighting that increased LT-HSC apoptosis reduced differentiation into ST-HSCs, ultimately lowering quiescent ST-HSC numbers. Overall, the PRCC analysis demonstrates the expected effects and sensitivity of various parameters on LT-HSC and ST-HSC populations, suggesting robustness in the model.

### Numerical Exploratory analysis of the spatial component of the model

The migration and spatial dynamics of HSCs remain poorly understood due to technical challenges in long-term in vivo visualization (Johansson et al., 2024; MacLean et al., 2017; Upadhaya et al., 2020). While HSC motility has been observed within the bone marrow and potentially toward the spleen under stress, direct evidence is limited. It is established, however, that HSCs exhibit great motility under homeostatic conditions and stress-induced scenarios.

In this work, our numerical exploratory analysis revealed gradients in cell distribution influenced by theoretical microenvironmental conditions, following a slow motion of cells, set to 600 µm/time-step, attributed to Brownian diffusion. While this motion is slow compared to other studies such as Upadhaya et al., 2020, where they showed mobility of 0.15 μm/min, implying a faster displacement per week than our simulated conditions, it is important to note that they observed HSC displacement over a period of only six hours. Therefore, no direct evidence supports the assumption that HSCs cover the same spatial distances within a seven-day timeframe.

In our results, when *aQ* and *aD* were set to 0.25, both active LT-HSCs and ST-HSCs exhibited a gradient towards the left side of Ly, with no clear preference for localization at the bottom or top of the Ly axis, due to the absence of changes in the cell cycle dynamics of LT-HSCs and ST-HSCs during the homeostatic simulation. Conversely, quiescent LT-HSCs tended to form a gradient toward the right side of Lx, near x=Lx, as time advanced, while quiescent ST-HSCs remained uniformly distributed in the spatial domain with an increase in numbers over time (time steps) consistent with the homeostatic configuration. The clear distinction in the location of both active and quiescent LT-HSCs and ST-HSCs reflects a theoretical heterogeneous spatial distribution and domain within the 2-D bone marrow niche, where quiescent cells tend to remain in G0 state in hypoxic regions typically localized perivascularly near sinusoidal vessels and arterioles (Acar et al., 2015; Chen et al., 2022; Kunisaki et al., 2013), while proliferating and differentiating HSCs exhibit more dynamic movement due to interactions with cytokines and chemokines (Baldridge et al., 2010; Essers et al., 2009; Jahandideh et al., 2020). These findings provide theoretical insights into how the bone marrow niche shapes HSC dynamics.

In addition, our exploratory analysis of when *aQ and aD* were set up to 0.5 and 0.75 independently, led to the assumption of the absence of a pronounced gradient of hypoxic regions within the 2-D spatial domain resulting in cell behavior consistent with a relatively homogeneous environment. Under these conditions, both quiescent and active LT-HSCs and ST-HSCs overlapped spatially, maintaining slow division rates. This underscores the critical role of localized hypoxia in shaping niche heterogeneity and HSC behavior.

Overall, our spatial model provides a versatile framework for simulating HSC dynamics within a theoretical 2-D bone marrow niche. By incorporating diffusion-based motility and spatial gradients, the model captures the interplay of microenvironmental factors regulating LT-HSC and ST-HSC behavior. These simulations offer valuable insights into the spatial regulation of HSC dynamics under homeostatic conditions, bridging gaps in experimental observations and advancing our understanding of the bone marrow niche.

## Materials and Methods

### Dynamical Stochastic Model Implementation

A computational framework based on single-cell stochastic behavior was developed to study the dynamics of LT-HSCs and ST-HSCs using both experimental and theoretical information.

The model incorporates the stepwise differentiation process, previously established by the Weisman group (Akashi et al., 2000; Kondo et al., 1997; Morrison et al., 1997), which outlines a hierarchical progression from LT-HSCs to ST-HSCs and then to MPPs. Our computational model employs latent stochastic processes, enabling the estimation of transition rates between cellular states and the changes in population composition over time. Additionally, it helps identify specific signatures associated with different states. These transitions are described by a non-homogeneous Markov process.

The model assumes that LT-HSCs and ST-HSCs undergo transitions between distinct cellular states, *S*(*t*), governed by specific transition probabilities. Mathematical S(t) can be represented as:

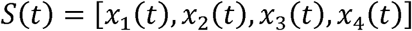

where each element represents a specific aspect of a cell state or cell fate.

- *x*_1_(*t*): This variable represents the **differentiation stage** of the cell. It takes on one of three values corresponding to different stages or cell fates: LT-HSC, ST-HSC, or MPP.
- *x*_2_(*t*): This variable captures the **cell cycle state**. It is a four-dimensional vector containing: mean cell division time, time since the last division, and the number of divisions, and whether the cell is active or inactive.
- *x*_3_(*t*): This variable captures the cell’s **quiescent state**. It is binary (0 or 1) indicating whether the cell is currently quiescent (not actively dividing) or dividing.
- *x*_4_(*t*): This captures the **cell apoptosis state**. It is a binary value (0 or 1) indicating whether the cell is undergoing programmed cell death (apoptosis).

### Model Assumptions

#### Stochasticity

Deterministic models, particularly those relying on ordinary differential equations (ODEs), can effectively depict the average behavior of HSC populations. However, they cannot capture the inherent variability observed in HSC dynamics. To overcome this limitation and provide a more accurate representation, our computational model uses stochastic simulations. These simulations incorporate randomness in two distinct mechanisms. The first mechanism involves the probabilities assigned to different cell fate options, such as differentiation, self-renewal, quiescence, or apoptosis. The second mechanism characterizes the variability in the duration of the cell cycle for each cell, modeled by Gaussian distribution functions.

#### Quiescence

HSCs reside in the bone marrow, where they predominantly remain in a relatively inactive, quiescent state known as the G0 phase (Calvi & Link, 2015; Göttgens, 2015; Nakamura-Ishizu et al., 2014). Despite their dormancy, HSCs possess a remarkable capacity for proliferation and differentiation. Previous studies under steady-state conditions have indicated that approximately 65% of HSCs are quiescent in specific regions of the bone marrow (Bradford et al., 1997; Cheshier et al., 1999). However, HSCs are highly dynamic and can exit the quiescent state, entering the G1 phase, to meet the system’s needs, causing fluctuations in the percentage of quiescent cells at any given time.

In our model, a predefined number of LT-HSCs will be evaluated for quiescence using the probability *PQA* to determine how many will undergo division. Those LT-HSCs that enter the cell cycle and divide will create ST-HSCs if the conditions are met. These ST-HSCs will then undergo a process of quiescence based on the probability *PQB*. Both processes are independent of each other.

To simplify the model, it is assumed that the quiescent state is irreversible; thus, cells that leave the G0 state cannot return to it during the simulation. Although quiescent cells do not actively participate in stem cell proliferation or differentiation, they remain susceptible to cell death with probabilities *PAQA* and *PAQB* for LT-HSCs and ST-HSCs respectively.

#### Cell Cycle

The divisional history of a cell is captured by its state variable, *x*_2_(*t*). This variable is a four-dimensional vector whose components are:

- ***Time since the last division***: This value is tracked through a separate counter within the code and reflects the elapsed time since the cell’s last division.
- ***Mean cell division time*:** This value represents the average time it takes for a cell, in its current state, to complete a full cell cycle. It is assigned to each cell independently from other cells, drawn from a Gaussian distribution with probability density function (PDF) described by the following equation:

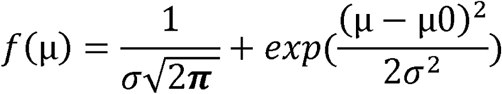

where µ represents the cell division time for a specific cell, µ_O_ is the population’s mean cell division time. In the code this has been denoted *meanCCA* and *meanCCB* for LT-HSCs and ST-HSCs, respectively. Finally, the standard deviation, *σ*, captures the variability in these division times across the population. In the model, we have denoted the standard deviations as *stdCCA* and *stdCCB*.

- ***Number of divisions***: The model incorporates a maximum number of divisions for LT-HSCs and ST-HSCs, representing the cumulative divisional history of these stem cells throughout their lifespan during active cell cycling. This variable reflects findings by Bernitz et al., 2016, who reported that slow-cycling HSCs can track their division history. Each cell has a limited number of divisions before entering a state of dormancy (reduced activity) associated with age-related phenotypic changes. Leveraging the model’s stochasticity for individual cells, this adjustable parameter allows us to explore how divisional history influences different outcomes in our simulations.
- ***Active/inactive cell***. The model incorporates the concept of inactivity once the cells (LT-HSCs and ST-HSCs) have reached a specific number of divisions, which is determined by the maximum number of divisions (*divA* and *divB*) threshold. In addition, active cells (LT-HSCs and ST-HSCs) are the cells that actively participate in the simulation.

#### Apoptosis

Apoptosis, a natural form of programmed cell death, plays a critical role in maintaining a healthy balance within cell populations (Elmore, 2007; Norbury & Hickson, 2001). This process, represented in the model by *PAA* and *PAB*, eliminates HSCs at specific probabilities depending on their type; LT-HSCs and ST-HSCs are assigned different turnover probabilities through *PAA* and *PAB*, respectively.

Previous studies have reported varying rates of HSC turnover. For instance, Kiel et al., 2007, found an approximate daily turnover rate of 6%, while Foudi et al., 2009, observed a slower pace ranging from 0.8% to 1.8% per day. This discrepancy could be due to the heterogeneity in the cell population and therefore it highlights the importance of using a flexible range for *PAA* and *PAB* in the model. By allowing these probabilities to vary, the simulations can explore a wider range of outcomes.

#### HSCs fate decisions

LT-HSCs that are not quiescent can either undergo self-renewal or differentiate during the simulation. Self-renewal can occur through symmetrical proliferation (creating two LT-HSC daughter cells) with probability *P1A* or asymmetrical proliferation (producing one LT-HSC and one ST-HSC) with probability *P2A*. Alternatively, differentiation can occur via direct differentiation (yielding one ST-HSC) with probability *P3A* or symmetrical differentiation (resulting in two ST-HSCs) with probability *P4A*. These types of divisions have been documented in experimental and mathematical studies of HSCs (Bernitz et al., 2016; Kawahigashi et al., 2024; Radtke et al., 2023).

ST-HSCs, derived from LT-HSCs, follow similar decision-making processes during the simulation. Both proliferative and quiescent ST-HSCs are susceptible to death with a probability *PAB*. Unlike LT-HSCs, ST-HSCs are known for more frequent cell division (H. W. Yang et al., 2020). Proliferative ST-HSCs contribute to their pool expansion through symmetrical proliferation (creating two ST-HSC daughter cells) with probability *P1B* and asymmetrical proliferation (producing one ST-HSC and one MPP) with probability *P2B*. Additionally, ST-HSCs can differentiate via asymmetrical differentiation (yielding one MPP) with probability *P3B* and symmetrical differentiation (yielding two MPPs) with probability *P4B*. A detailed description of the parameters can be found in Supplementary Table 1, while Figure 1 provides a visual representation of the overall process. Notably, for both LT-HSCs and ST-HSCs, the sum of all division probabilities equals 1, reflecting the framework of Markov processes.

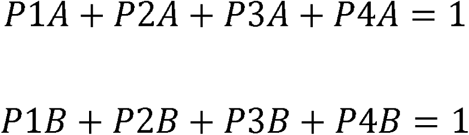

#### Spatial Stochastic Model Implementation

The spatial component of the model simulates the motion of LT-HSCs and ST-HSCs within a two-dimensional spatial domain bounded by L_x_ and L_y_, extending from x = 0 to x = L_x_, and from y = 0 to y = L_y_. The model records the position of each one of the quiescent cells and active LT-HSCs and ST-HSCs during the simulation within the spatial domain. The change in quiescent in space is determined by *aQ* and the change division rate in space is determined by *aD*. The spatial motion of both quiescent and active cells is determined by the Euler-Maruyama method (Kloeden & Platen, 1992), a first-order numerical method for approximating solutions to stochastic differential equations. The spatial model is governed by the following parameters:

- *dtBrow*: This variable determines the time discretization used for simulating Brownian motion.
- *Df*: This variable determines the diffusion coefficient of cells.
- *aQ*: This variable determines the gradient change in space of Quiescent cells
- *aD*: This variable determines the gradient change in space of mean division time.

##### Spatial gradients

Gradients of model parameters are implemented as linear functions of position. A parameter value (*P_β_*) at a specific location is calculated as:

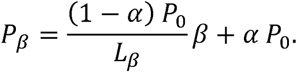

Here, *P*_0_ represents the parameter’s baseline value (i.e., its value if it were constant across the domain), *β* denotes the spatial coordinate (x or y), *L_β_* is the length of the domain along the *β* axis, and *α* determines the extent of the parameter’s decrease from one end of the domain to the other.

We consider spatial variation of the probability of quiescence (*pAQ or pBQ*) along the x-axis and of the mean cell division time (*meanCCA or meanCCB*) along the y-axis, resulting in the following equations for these spatially varying parameters:

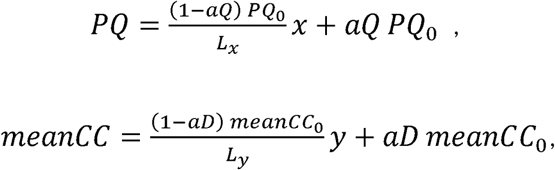

where *PQ*_0_ and *meanCC*_0_ are the values given in Table 1.

While our dynamical model does not explicitly simulate bone marrow niche interactions or external factors influencing HSC dynamics, we can indirectly infer their effects by analyzing the impact of model parameters and their resulting outputs. For instance, a lower probability of symmetrical proliferation could reflect impaired self-renewal capacity due to high concentrations of IFN-γ in the bone marrow under specific conditions (Baldridge et al., 2010). Thus, interpreting the model’s output is crucial for understanding how intrinsic cellular dynamics are modulated by the complex niche environment. The model’s spatial component allows us to explore the movement of quiescent and active LT-HSCs and ST-HSCs within a 2D bone marrow niche, based on the assumption that HSCs can migrate under both homeostatic and stress conditions (Johansson et al., 2024; Miao & Pereira, 2020). Although our model uses Brownian motion as a baseline for HSC movement, in vivo studies demonstrate more complex migration patterns during infection, including increased heterogeneity and sustained migration (MacLean et al., 2017). Our model explores how spatial gradients may contribute to these deviations from Brownian motion. We acknowledge that other migration mechanisms, such as directed or persistent random walks, may also play a significant role and will be the subject of future work.

#### Model Limitations

Developing a comprehensive mathematical model for a complex biological system, such as hematopoietic stem cell (HSC) dynamics, presents inherent challenges. While our current model offers a powerful framework to study HSC dynamics from a stochastic and spatial perspective, it can be further refined to incorporate additional biological complexities. One potential area for improvement lies in integrating a multi-scale approach. This could involve explicitly modeling the cell cycle phases (G1, S, G2/M) that HSCs undergo before division. Currently, our model treats proliferation as a single event but incorporating these phases could provide a better understanding of cell cycle regulation and its impact on HSC behavior. Furthermore, our model assumes an irreversible transition from the active cycling pool to a quiescent state. However, it is known that quiescence is an irreversible process, where cells can re-enter the G0 state in response to stimuli. Some studies suggest that under specific conditions, HSCs might reactivate from quiescence. Accounting for this potential reversibility within the model framework could enhance its biological realism and allow for the exploration of scenarios involving HSC exhaustion or rejuvenation therapies. On the other hand, we proposed a model that describes the stochastic motion of both quiescent and active LT-HSCs and ST-HSCs within a two-dimensional space niche. While we aim to shed light on the movement and positioning of active LT-HSCs and ST-HSCs alongside their quiescent counterparts, it’s essential to recognize the complexity of their biology; it extends beyond just these two dimensions. Future developments will involve enhancing the model to incorporate additional biological factors, such as the cell cycle, and advancing toward a three-dimensional representation of the bone marrow niche environment.

#### Parameter Sensitivity analysis

To examine robustness and the relationship between model parameters and outputs, we employed the Latin Hypercube Sampling (LHS) technique in combination with Partial Rank Correlation Coefficient (PRCC) analysis. We assumed that each uncertain parameter follows a uniform distribution within a specified range. The LHS method was implemented by segmenting the value ranges of each parameter into equally probable intervals (inputs), ensuring a comprehensive exploration of parameter space (Gomero, 2012).

LHS has a minimum required sample size (*n*) which is given by *n*≥*k*+1 *or n*≥(4/3)*k* where *k* is the number of parameters included in the LHS (Blower & Dowlatabadi, 1994). We generated 10,337 parameter combinations from the chosen parameter distributions, with model outputs evaluated at time step 300 to capture the full range of model behavior. The area under the curve (AUC) was quantified using the trapezoidal integration method.

PRCC then was calculated for each of the following parameters: *P1A, P2A, P3A, P1B, P2B, P3B, PQA, PQB, PAA, PAB, meanCCA, stdCCA, meanCCB, stdCCB, divA, and divB* (see Supplementary Table 1), and the outcome variable (the total number of active, quiescent and inactive LT-HSCs and ST-HSCs). The **s**ign of the PRCC indicates whether changes in a parameter have a positive or negative effect on the corresponding output variable. Additionally, statistical z-scores and their corresponding p-values were calculated for each parameter and correlation coefficient to assess statistical significance during the LHS/PRCC analysis.

The z score for the Spearman rank correlation coefficient ρ was computed as:

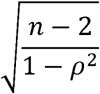

where *n* is the number of samples. The p-values were computed as:

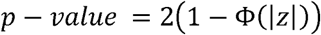

where Φ(∣z∣) is the cumulative distribution function of the standard normal distribution evaluated at the absolute value of the z score, |z|.

PRCC analyses are displayed as bar plots from -1 to 1 for each parameter and their respective output. The parameters that show an * on top are the ones with a statistically significant correlation (p<0.05).

#### Numerical Exploratory analysis of the spatial component of the model

In this section, we conducted a numerical exploratory analysis to investigate the theoretical spatial dynamics of LT-HSCs and ST-HSCs in a two-dimensional niche mimicking the bone marrow niche. Using the homeostatic configuration fit to experimental data from Säwen et al., we explored cell dynamics of quiescent and active LT-HSCs and ST-HSCs at time steps 20, 40, 60, and 80 by modifying the *aQ* (*PQA/PQB*: gradient of becoming quiescent) and *aD* (*meanCCA/meanCCB*: gradient of mean proliferation rate), as well as fixing the Brownian diffusion coefficient to 600 µm/time-step. The parameters *aQ* and *aD* introduce spatial heterogeneity by modulating cell properties based on their positions within the domain. For instance, cells located near one end of the domain (x=0) may exhibit a lower probability of becoming quiescence, while those near the opposite end (x=Lx) may display a higher probability of becoming quiescence. Similarly, cells at the bottom of the domain (y=0) might exhibit a higher division rate, whereas those at the top (y=Ly) might divide more slowly.

The *aQ* and *aD* gradients were uniformly modified as follows below, and those cases were studied independently:

- *aQ* =0.25, *aD* =0.25
- *aQ* =0.5, *aD* =0.5
- *aQ* =0.75, *aD* =0.75

## Acknowledgements

The authors thank the University of South Carolina’s Hyperion High-Performance Computing cluster for all the resources provided to perform this work.

## Competing interests

Authors declare no competing interest

## Funding

We acknowledge the support from Grant Nos. NIH R03DK124738, NIH K01DK104974, NIH P20GM109091, and USC ASPIRE 130100-24-68351. to K.L.K. and Grant No. NSF-DMS-1751339 to P.A.V.

## Data and Resource availability

The data and codes that support the findings of this study are available from the corresponding authors upon reasonable request.

---

**Table 1.Parameters used to describe independent homeostatic LT-HSCs and ST-HSCs dynamics according to Busch et al. and Säwen et al.** Both simulations were performed for up to 80 weeks. Both simulations did not take into consideration spatial changes to fit our model to mouse homeostatic conditions according to the references. The 20 dynamical parameters are listed with their corresponding value for each independent dataset.

**Table 2. Parameters used to describe all the apoptotic-inflammatory stress conditions on LT-HSCs and ST-HSCs dynamics** Both simulations were performed up to 80 weeks. Table displays the parameter combination used to fit and mimic apoptosis-related scenarios on LT-HSCs and ST-HSCs dynamics. All these simulations did not take into consideration spatial changes to fit our model to mouse homeostatic conditions according to Read et al. The 20 dynamical parameters are listed with their corresponding value for each independent scenario.

**Table 3. Parameters used to model theoretical numerical scenarios of the spatial distribution of LT-HSCs and ST-HSCs.** Table displays the parameter combination used to model spatial gradients and distribution of LT-HSCs and ST-HSCs using different *aQ* and *aD* combinations, at time steps 20, 40, 60, and 80. Homeostatic configuration fit to Säwen et al. is used to investigate spatial motion. The 26 dynamical and spatial parameters are listed with their corresponding value.

## Notes

### Competing Interest Statement

The authors have declared no competing interest.

