## Supplemental Table 1 for "STOCHASTIC MODELING OF HEMATOPOIETIC STEM CELL DYNAMICS"

**Supplementary Table 1. Model parameters describing LT-HSCs and ST-HSCs dynamics in the stochastic and spatial model:** The 28 parameters are listed in this table with their corresponding description. There are 10 parameters that control the dynamics of LT-HSCs and 10 parameters for the dynamics of ST-HSCs. The last 6 parameters correspond to spatial component of the model

| **Parameter** | **Description** |
|---|---|
| *t* | Time frame of the simulation. |
| $N_A$ and $N_B$ | Initial number of LT-HSCs and ST-HSCs. |
| PQA | Quiescence probability for LT-HSCs |
| PQB | Quiescence probability for ST-HSCs |
| P1A | LT-HSCs symmetrical proliferation |
| P2A | LT-HSCs asymmetrical proliferation |
| P3A | LT-HSCs direct differentiation |
| P4A | LT-HSCs symmetrical differentiation |
| P1B | ST-HSCs symmetrical proliferation |
| P2B | ST-HSCs asymmetrical proliferation |
| P3B | ST-HSCs direct differentiation |
| P4B | ST-HSCs symmetrical differentiation |
| PAA | LT-HSCs apoptosis |
| PAB | ST-HSCs apoptosis |
| PAQA | Quiescent LT-HSCs apoptosis |
| PAQB | Quiescent ST-HSCs apoptosis. |

| | |
|---|---|
| meanCCA | LT-HSCs mean cell division time. |
| meanCCB | ST-HSCs mean cell division time. |
| stdCCA | Standard deviation of the LT-HSCs cell cycle. |
| stdCCB | Standard deviation of the ST-HSCs cell cycle. |
| divA | Maximum number of divisions for active LT-HSC. |
| divB | Maximum number of divisions for active ST-HSC. |
| Lx | First spatial dimension size. |
| Ly | Second spatial dimension size. |
| dtBrow | Time discretization for Brownian motion |
| Df | Diffusion coefficient |
| aQ | Gradient change of quiescent cells |
| aD | Gradient change of mean division time |
